# Examining Action Potential Waveform Diversity in Neuronal Populations of Midbrain Organoid Models

**DOI:** 10.64898/2026.06.19.733318

**Authors:** Juraj Ondriš, Anna-Sophie Zimmermann, Daniele Ferrante, Jens Christian Schwamborn

**Affiliations:** Cognitive and Systems neuroscience, Swammerdam Institute for Life Sciences, University of Amsterdam, Amsterdam, The Netherlands; Developmental and Cellular Biology, Luxembourg Centre for Systems Biomedicine, University of Luxembourg, Belvaux, Luxembourg

**Keywords:** Parkinson’s disease, midbrain organoid, clustering, extracellular electrophysiology, multi-electrode array, spike-sorting

## Abstract

Over the last decade, pre-clinical research has witnessed the advancement of human induced pluripotent stem-cell derived 3D brain organoid models and their differentiation into specific brain regions. In the realm of Parkinson’s disease research, development of midbrain-specific organoids has enabled studies of this neurodegenerative disorder in patient derived 3D organoid models that attempt to recapitulate the human brain complexity. Within this line of research, neural functionality of the organoid models is established through electrophysiology. As a novel methodological approach, this study aimed to establish whether clustering of electrophysiological activity originating from midbrain organoids would aid in identifying different types of action-potential waveforms exhibited by neurons within the organoid model. Long-term extracellular electrophysiological recordings were conducted by use of a multi-electrode array device. The local field potential signal was spike-sorted, and the extracted putative neuron units were clustered into groups of spike waveform profiles. After establishing this methodological analysis pipeline, the clusters of waveform types were further analyzed in terms of electrophysiology. Results revealed that the clustering approach was successful at identifying three types of spike waveforms categories. Furthermore, it was proposed that one spike waveform profile potentially originated from dopaminergic neurons, which were one on the neural cells populating the organoid models. Overall, this study has successfully established a new methodological clustering approach to analyze electrophysiological data recoded from 3D organoid models in the context of Parkinson’s disease modelling and organoid model development research.

## 1 Introduction

Throughout the course of the recent decade, groundbreaking developments have improved the available methods and models to study neurodegenerative disorders. More specifically, the generation of human induced pluripotent stem cells (iPSCs) from dermal fibroblasts(Takahashi et al. 2007) opened a plethora of new ideas into pre-clinical research on this subject. Formerly, research was conducted with limited tissue specimens from patients that involved complicated and invasive methods of extraction(H. Braak et al. 2000; Heiko Braak et al. 2003). Other research focused on animal models to investigate human neurodegenerative disorders(Betarbet et al. 2002; Dawson et al. 2002), which eventually led to the inescapable impasse of interpreting the relevance of cross-species comparisons, given complex biological distinctions between humans and animals. The advancement to generate human iPSCs has not only made it easier to collect samples, but also brought research closer to the molecular nature of diseases. In itself, the iPSC technology still experiences new developments. From fibroblast derived iPSCs, to monolayer two-dimensional (2D) cell culture systems of differentiated neuronal tissue(Friling et al. 2009), to three-dimensional (3D) organoid tissue structures(Monzel et al. 2017). Brain organoid models attempt to recreate localized brain regions with specifically differentiated cell populations. These novel innovations facilitate advanced *in-vitro* disease modelling, which holds clinical potential for future individualized patient therapies.

One area of research where such methods are applicable is targeted at studying Parkinson’s Disease (PD). This neurodegenerative disorder leads to a cascade of motor-related symptoms (i.e., involuntary tremors at rest, bradykinesia, limb rigidity, gait and balance problems), but is also known to show other non-motor manifestations like cognitive impairments or sensory disturbances(Olanow et al. 2009). PD is the most common movement disorder, with its recently assessed global prevalence estimated to 8.5 million individuals in 2019(World Health Organization: WHO. 2023). Given that PD related neurodegeneration predominantly affects dopaminergic neurons (DNs) in the substantia nigra pars compacta(Inamdar et al. 2007), recent studies set out to generate organoids that would encompass this neuronal population from the midbrain region. For instance, one study devised a human midbrain-specific organoid (MO) model from neuroepithelial stem cells aimed to directly model PD(Monzel et al. 2017). Apart from having spatially organized DNs, the organoids showed various populations of astroglia, oligodendrocytes, and neurons. They further observed electrophysiological activity, synaptic communication, and myelination of processes. Additionally, single-cell RNA sequencing of the MO identified the following neuronal cell types: young neurons, mature DNs, non-DNs, and neuroblasts (neuron-like progenitor cells)(Zagare et al. 2022). These findings confirm that the currently examined MO model encompasses DNs as well as other neuron types.

Advancing models is crucial for expanding scopes of research, providing more representative models of the true nature of the object of study. Nevertheless, further investigation and refinement of analysis methods is also necessary. Since PD is characterized by neurodegeneration of DNs, electrophysiological assessment of brain organoid models is of great interest. It can provide a form of evaluation of neural functionality and enables comparisons of functionality between disease models. For example, electrophysiological activity may be studied through direct sample stimulation with trial-based designs, drug treatments to modulate specific ionic-channel movements, or by measuring spontaneous activity levels and conducting between-group comparisons of electrophysiological measures such as firing rate, connectivity, oscillatory dynamics, or bursting of neurons(Shin et al. 2021; Smits et al. 2020; Sharf et al. 2022). Considering that PD predominantly exhibits its neurodegenerative impact on DNs, investigating electrophysiological activity of DNs in isolation is relevant. Focusing on iPSC derived samples, whole-cell *in-vitro* patch-clamping is one way to directly record DNs(Bardy et al. 2016). Although this ensures direct intracellular electrophysiological readings from DNs, some limitations arise. The DNs are generated in monolayer 2D cultures. Compared to more complex 3D organoid models, 2D cultures of neurons are less complex in terms of recapitulating brain physiology. Moreover, conducting patch-clamp recordings of specific neurons in 3D organoids models is extensively difficult, as the tissue is hard to navigate, and specific cells are embraided within other neuronal cells that obstruct the way. Another method to directly record DNs is via two-photon calcium imaging, as demonstrated by a previous study that stained midbrain DNs differentiated from iPSCs, enabling calcium imaging of specific neurons as a method of recording electrophysiological activity(Hartfield et al. 2014). Despite that, this method also revolves around monolayer cultures of DNs in isolation. With regards to calcium imaging of more advanced 3D organoid models, selectively measuring the ionic-channel influx of calcium in specific neuron types is more difficult to implement, requiring a more complex experimental design.

A practical way to measure electrophysiological activity of 3D organoid models is via multi-electrode arrays (MEA). This device allows recordings of surface local field potentials (LFP) in *ex-vivo* sections of brain tissue(Zoladz 2013), or *in-vitro* 2D cultured cells and 3D organoid models(Pelkonen et al. 2021). MEAs capture LFPs through various grid layouts of electrodes in proximity of the recorded tissue. This method is favorable for obtaining electrophysiological activity of 3D organoid models, as its application is fast and efficient. As shown by previous research(Monzel et al. 2017; SabateLJSoler et al. 2022), this non-invasive long-term recording setup yields electrophysiological data of cells on the outer surface of organoid models. However, with this data at hand it remains challenging to attribute signals to specific neurons and classify them into biological categories of neurons. It would therefore be advantageous to unravel the signal in the form of spike/action-potential (AP) waveforms and distinguish them into different classes, which could potentially support inferences regarding the biological identity of neurons that generated them. This may be relevant for isolating electrophysiological activity of DNs in 3D organoid models, which are subject to neuroinflammatory neurodegeneration in PD(Shabab et al. 2017). This new methodological approach could be beneficial by enabling evaluations of DN functionality between disease models.

To interpret potential types of neurons recorded by the MEA, it is firstly necessary to determine a computational approach of clustering neuron AP waveforms. One such clustering method was previously applied to extracellular recordings of neural activity in the cat primary visual cortex (V1)(Sun et al. 2021). First, activity was spike-sorted to extract neuronal spike waveforms. These waveforms were classified into five separate categories based on template metrics of waveform shapes. The categories were labelled based on characteristics of the waveforms present in each group, namely: regular-spiking, fast-spiking, triphasic-spiking, compound-spiking, and positive-spiking waveforms. This categorization enabled comparisons of electrophysiological metrics such as: firing rate, bursting rate, or response latency between waveform categories. This method was also deployed in a previous study, to cluster extracellularly recorded spike waveforms from the wallaby V1(Jung et al. 2023). Results highlighted the identification of similar cluster categories to the abovementioned study. Despite that, these former clustering analyses were applied to data from animal V1, meaning the species and brain region was different in comparison to currently investigated MOs. Even though cell types remain stable across species, there are differences in cell development, proliferation and structural organization between human midbrain neurons, differentiated from iPSCs, and mouse midbrain neurons(La Manno et al. 2016). Nonetheless, more prominent differences exist between V1 and midbrain cortical regions. The types of neurons residing in human MOs(Zagare et al. 2022) are heterogenous with respect to neurons identified in the mouse V1(Gouwens et al. 2019).

Given the contrast between MOs and the mouse and wallaby V1, the implemented clustering procedure was adapted to accommodate various types of neuron spike waveforms observed in electrophysiological recordings. Considering this context, the present study aimed to investigate whether clustering of spontaneous electrophysiological activity could assist in identifying different types of AP waveforms exhibited by neurons within MOs. Establishing such a computational pipeline was hypothesized to uphold the possibility of classifying spike waveform types from the observed LFP activity. Secondly, it was hypothesized that such waveform categories of responses could theoretically be linked to known neuronal populations within the recoded organoid models.

## 2 Materials and Methods

### 2.1 3D Cell Culture

#### 2.1.1 Midbrain Organoids

The generation and culturing of human MOs was conducted as described in previous literature(Nickels et al. 2020; Smits et al. 2019). Firstly, ventral-midbrain identity pre-patterned neuroepithelial stem cells (vNESCs) were cultured in 6-well plates that were pre-coated with Matrigel (BD Bioscience). The vNESCs were maintained using N2B27 medium supplemented with 0.2mM Ascorbic acid (Sigma, A4544), 3μM CHIR (Axon Medchem, CT 99021), 0.5μM Smoothened Agonist, SAG (Merk Millipore, 566660), 2.5μM SB-431542 (Abcam, ab120163), and 0.1μM LDN-193189 (Sigma, SML0559). The N2B27 medium comprised of DMEM-F12 (Invitrogen)/Neurobasal (Invitrogen) 1:1 with 1:200 N2 (Invitrogen), 1:100 B27 lacking Vitamin-A (Invitrogen), 1% L-glutamine, and 1% penicillin/streptomycin (Invitrogen)(Monzel et al. 2017).

After reaching a confluency of ∼80%, the cells were detached using Accutase (Invitrogen). For seeding (day-0), 6000 vNESCs were plated in each well of an ultra-low attachment 96-well plate (Corning), in the aforementioned medium. The plates were centrifuged (300xg for 3mins) to support formation of 3D spheroids. On day-2 the cells underwent the first pre-patterning process where SB-431542 and LDN-193189 were withdrawn from the supplemented N2B27 medium. Following this, on day-4 the concentration of CHIR was lowered to 0.7μM in the second pre-patterning process. On day-8, the spheroids were embedded onto MEA chips. Hereafter, the MOs were cultured in co-culture maturation medium, consisting of Advanced DMEM/F12 (Thermo Fisher, 12634010), 1×N2 (Thermo Fisher, 17502001), 1×Pen/Strep (Invitrogen, 15140122), 1×Glutamax (Thermo Fisher, 35050061), and 50μM 2-mercaptoethanol (Thermo Fisher, 31350-010). This medium was supplemented with 100ng/mL IL-34 (Peprotech, 200-34), 10ng/mL GM-CSF (Peprotech, 300-03), 10ng/mL BDNF, 10ng/mL GDNF, 10μM DAPT and 2.5ng/mL Activin A. The spheroids were kept under static conditions with regular media exchanges (every 7-days) supporting their differentiation. All cultures were kept in incubators with 5% CO_2_ at 37 °C with a relative humidity >95%.

#### 2.1.2 Plating Organoids onto MEA Chips

The embedding of MOs was implemented directly on MEA chips. To begin with, MEA chips were thoroughly cleaned by applying 1mL of 1%-Terg-a-zyme-solution enzyme detergent (10g/L in deionized water, Sigma-Aldrich) inside the chip culture chambers. The enzyme-covered surface was gently agitated with a paintbrush and placed onto a shaker for ∼3h. Subsequently, the detergent was removed, and culture chambers were washed three times with 1mL DPBS. The chips were left to dry in a ventilated biosafety cabinet before being enclosed with a tight seal cap, to provide a sterile environment for organoids to be cultured and recorded in.

After cleaning, the chips were coated. As primary coating, polyethyleneimine (PEI, Sigma-Aldrich) was diluted in sterile borate buffer to a 0.07% concentration. PEI was applied in 50µL droplets covering the chip electrodes, and then incubated for 60-minutes. The primary coating was then aspirated, the chip was washed with deionized water, and left to dry in a sterile environment for 60-minutes. As secondary coating, Laminin (0.04mg/mL of the co-culture medium, Sigma-Aldrich) was administered in 50µL droplets spreading across the chip center. After 60-minutes, Laminin was aspirated leaving a thin layer of coating on the electrodes surface. Following the coating procedure, organoids were plated by gently transferring the cultured spheroids to the center of the electrodes. Excess fluids were aspired and the organoids were left to properly attach to the coatings for ∼2-minutes in an incubator. Thereafter, a 50µL droplet of GelTrex (Thermo-Fisher) was deposited on top of the plated organoids and incubated for 5-minutes to support attachment of the sample at the chip center. Finally, culture chambers were filled with 1000µL of the co-culture maturation medium and sealed with caps before being transferred to the incubator. Medium exchanges were carried out once a week, at least one day prior to scheduled recording days.

### 2.2 Electrophysiology

#### 2.2.1 MEA Recordings

This experiment was directed at analyzing electrophysiological activity of MOs generated from two different cell lines (demographics in Table 1). Due to project-related constraints in resources and time, we examined one MO model per cell line. Spontaneous electrophysiological activity was recorded employing the USB-MEA256 system (MCS, Germany) setup. This device utilizes Silicon nitride MEA chips for recording extracellular LFP. We used MEA chips with 256 circular 30µm wide titanium nitride electrodes, spaced in a square layout at a 100µm distance from neighboring channels (256MEA100/30iR-ITO-pr-T, MCS, Germany). The organoids, attached and cultured on the chips, were placed into the signal-amplifier recording slot on top of a heating element, connected to a temperature controller device (TCX-Control V1.3.4, MCS, Germany), maintaining a stable temperature of the samples at 37±0.5°C. After thoroughly closing the hatch to ensure good chip-to-device contact, a custom-made Faraday cage was placed on top of the hatch to shield recordings from outside electrical noise.

**Table 1.** Participant demographics and genetic specifications of cell lines utilized in the study. The acronym consists of a patient identifier ‘IM5’ followed by a specification of the cell line gene type (mutant – Mut, gene-corrected – GC). The LRRK2-G2019S mutation is one of the most common genetic determinants of PD. This information was directly derived from the cell-line database.

| Cell Line Demographics |  |  |  |  |  |
| --- | --- | --- | --- | --- | --- |
| Database Number | Acronym | Genetic Mutation | Condition | Sex | Age |
| 209 | IM5 Mut | LRRK2-G2019S | PD | F | 81 |
| 212 | IM5 GC | LRRK2-G2019S | PD → WT | F | 81 |

The samples were kept in this setup for 5-minutes to stabilize organoids in the new environment following the transfer. During recordings, culture chambers were grounded via four internal reference electrodes connected to ground. Signals were pre-amplified, digitalized, and recorded at 25kHz sampling rate in the MC_Rack software (MC_Rack V4.6.2, MCS, Germany). To minimize potential data-loss, the signal from all 252 electrodes was recorded in raw analogue format in 10-minute segments and stored for later pre-processing and filtering. Electrophysiological recordings were conducted longitudinally every five days, beginning on day-15 from organoid seeding, continuing until day-70. Recordings were performed both in the morning and afternoon to increase the amount of collected data.

#### 2.2.2 Signal Pre-processing

The pre-amplified raw analogue signal was pre-processed in MC_Rack software. Data was filtered with a high-pass 200Hz and low-pass 2000Hz filter. Following this, active electrode channels were selected to reduce file size for later analyses and increase the spike-sorting algorithm output quality. This selection was conducted by applying a spike detector threshold at 6 standard deviations (SD) of the filtered signal. A channel was manually labelled active if it presented a minimum of ten spikes across the recording time (1spike/min). Furthermore, channels showing fluctuating noise over time were excluded. After re-recording selected active channels, the data was converted to an appropriate file format for further analyses.

#### 2.2.3 Spiking Analysis

To establish electrophysiological activity and waveforms of individual putative neurons, the dataset was spike-sorted with Spikeinterface(Buccino et al. 2020). This is a Python(Anaconda Inc. 2020) based package that offers scripts for: universal data importing, probe generating, preprocessing, spike-sorting, postprocessing, visualizing, and exporting data. The following steps were encompassed in a script coded via Jupyter(Kluyver et al. 2016) to individually spike-sort 10-minute LFP recordings.

Firstly, re-recorded raw analogue data was imported and a custom-generated probe resembling the MEA chip channel layout was attached to the recording. Then, data was scaled, filtered with a high-pass 200Hz and low-pass 2000Hz filter, and referenced to the common median per channel. Pre-processed channel signals were spike-sorted with Tridesclous(Garcia and Pouzat 2016), a template-matching algorithm that automatically identifies putative units from extracellular activity. Spikes were detected using a threshold method of 6SD, and the extracted waveforms were then clustered into individual putative neuron units. This analysis also conducted spatial whitening, thereby detecting correlations across channels in an attempt to eliminate replicates of units, noise and artefacts. Consecutively, these sorting results were postprocessed with available functions to calculate template and quality metrics per detected unit. Employing these metrics, units were then automatically curated based on thresholds with the following variables: inter-spike-interval violation ratio < 0.2, firing rate > five spikes/minute, amplitude median > 10µV, and silhouette metric (goodness measure of PCA cluster representativeness) > 0.4. Units that failed to pass were labelled as multi-unit activity or noise and excluded from later analyses.

Automatically curated units were visually inspected in Phy(Phy 2020), a Python library for visualization of spike-sorted electrophysiological data. This library displays: spike amplitudes, autocorrelation functions, inter-spike-interval histograms, spiking patterns over recording channels/time, and spike waveforms with gaussian convolutions for visual assessment. This tool was used for checking automatic curation quality and tuning exclusion criteria. Finally, spike-sorting results and unit template/quality metrics were exported for further analyses.

### 2.3 Clustering and Statistical Analyses

#### 2.3.1 Threshold Clustering

The threshold clustering method was adapted from previous literature(Sun et al. 2021). This approach, based on waveform-shape metrics and thresholds, separates individual unit-waveforms into clusters resembling a homogenous waveform type. Adjustments made for this specific analysis regarded finetuning of previous clustering thresholds and employing new waveform shape metrics. The shape of each spike waveform was condensed into eight distinct feature measurements: amplitude, 1^st^ peak-to-trough ratio, 2^nd^ peak-to-trough ratio, spike duration, repolarization slope, recovery slope, and two novel metrics termed the ‘peak-to-trough ratio difference’ and ‘biphasic metric’. These measurements were collected from average waveforms of all recorded spikes per unit. Using spike-sorting postprocessing algorithms, the individual average waveform cutouts, unsigned amplitudes, spike durations, repolarization slopes, and recovery slopes were automatically determined prior to calculating the remaining waveform template metrics. A waveform baseline was established as the mean between magnitudes of the beginning and end point of each average waveform cutout to calculate the additional metrics.

To compute amplitudes, the absolute magnitude of waveform minimum and maximum peak was compared. The largest of the two was used to calculate the difference from baseline, representing the signed amplitude. The 1^st^ and 2^nd^ peak-to-trough ratios were determined by dividing the positive peak magnitude by the negative trough magnitude, using the first and second positive peaks, respectively. Further, to establish the ‘biphasic metric’, average unit waveform cutouts and waveform baselines were normalized between -1 and 1. The metric was then calculated as follows:

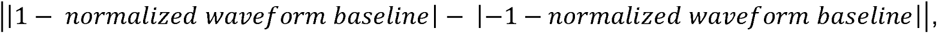

resulting with a feature that informs on the difference between the maximum peak and minimum trough. For an example waveform, a smaller measurement of this metric would suggest that the peak and trough had approximately equal unsigned amplitudes, indicating that the waveform probably resembled a biphasic spike. This feature was computed from normalized waveform baselines, as this standardizes the biphasic metric across all recorded unit waveforms. Finally, the ‘peak-to-trough ratio difference’ was deduced by computing the absolute difference between the 1^st^ and 2^nd^ peak-to-trough ratios.

After extracting waveform template metrics, unit waveforms were pooled together from all available recordings per organoid model. The pooled units were then clustered as follows. The first cluster consisted of units with a ‘biphasic metric’ < 0.55. The second cluster encompassed units with a signed amplitude > 0. The third cluster contained units with the ‘peak-to-trough ratio difference’ < 0.05. Remaining units were grouped to form the fourth cluster. These thresholds and metrics were chosen for the clustering analysis following trial and error piloting, selecting metrics and thresholds that yielded the most favoring clustering results. Other waveform shape features were thus omitted, to produce clusters more representative of the available waveforms. The clustering analysis, calculations of waveform shape metrics, and all further analyses were carried out in MATLAB(The MathWorks Inc. 2022).

#### 2.3.2 t-distributed stochastic neighbor embedding (t-SNE)

To visualize clustered groups of spike waveforms from a different perspective, t-SNE(van der Maaten and Hinton 2008; Yang et al. 2022) was performed. This is a dimensional reduction algorithm, which embeds waveforms into an n-dimensional representation, in this case, a 2D space. Each waveform in the t-NSE latent dimensions is represented as a single point-vector with coordinates. The position of the point and its distance to others is established based on its similarity to other points. In this case, the location of a unit waveform vector in this space would be determined by its similarity or dissimilarity to other waveforms. This analysis allows for a visual assessment of waveforms in the t-SNE representation, labelled according to their cluster, and allows inspection of how effectively the clusters group unit-waveforms based on their similarity. It may additionally be used to highlight cluster quality aspects, for example contamination by outliers that could belong to different clusters.

Input data for the t-SNE algorithm consisted of average voltage traces across time for each spike waveform (2D matrix, with units × timepoints). As the t-SNE computes similarity in terms of explained variance, it internally runs an initial principal component analysis before embedding waveforms in the latent space. Some input parameters required manual configuration during setup and were defined according to prior research(van der Maaten and Hinton 2008; Yang et al. 2022). The ‘initial PCA dimensions’ variable was set to 50, ‘t-SNE distance’ to cosine, ‘verbose’ to equal two, and ‘perplexity’ to 30. Finally, the number of t-SNE dimensions to embed waveforms in was configured to two, later enabling a straightforward 2D visualization for plotting results.

#### 2.3.3 K-means Clustering

To compare threshold clustering results to an automated clustering algorithm, the k-means clustering analysis was conducted, sourced from MATLAB(The MathWorks Inc. 2022). This algorithm iteratively assigns data points to a manually predetermined number of clusters, denoted as ‘k’. In this case, as results were compared to the threshold clustering approach, k was set to equal four. The input data consisted of x- and y-axis vectors from each unit-waveform in the 2D t-SNE representation. The algorithm began by determining cluster centers, termed centroids, using a heuristic approach adapted from previous research(Arthur and Vassilvitskii 2007). The algorithm then calculates all distances of points to cluster centroids, and assigns each to the closest centroid. The distance of points to their respective cluster was set to ‘cityblock’, defined as the sum of absolute differences, meaning centroids resembled the median of its point members. Finally, this process was repeated 50 times, as initialized cluster centroids differ each run. The run with lowest sum of point-to-centroid distances was selected for reporting. Resulting unit waveform cluster identity from the automatic k-means algorithm was then compared to the threshold clustering results.

#### 2.3.4 Statistical Analyses

Statistical between-group comparisons were implemented non-parametrically, as some data groups had insufficient sample sizes (n<15). Additionally, all statistical analyses tested comparisons between more than two groups. Therefore, the Kruskal Wallis test was chosen for testing statistical differences, with alpha set to 0.05. To test differences of means between individual groups, post-hoc multiple comparison analyses were conducted. The Dunn-Sidak alpha value correction method was utilized to protect from family-wise error rates resulting from multiple pairwise comparisons.

## 3 Results

### 3.1 Spike-sorting LFP Activity Revealed Various AP Waveforms Across Organoid Models

Prior to clustering waveform profiles, spike-sorting was conducted to isolate putative units and their associated APs from the LFP MEA recordings. Automatic curation of identified units formed the dataset of putative neuron AP waveforms, which were the primary focus of later analyses. Due to an insufficient number of units, the MO from the mutant line was excluded. Additionally, day-15 was excluded given a low sample size of detected units. Furthermore, observations indicated that organoids exhibited abnormal morphologies, signs of deterioration, or began detaching from MEA chips following day-45. Consequently, all data recorded beyond this timepoint was not considered for further analyses. The remaining dataset comprised of putative neuron units recorded from the gene-corrected MO in timepoints ranging from day-15 to day-45.

Examining individual waveforms of neuron APs highlighted varying types of waveform shapes throughout each recording. Fig. 1 depicts nine randomly chosen unit-waveforms from the MO recording on day-20. Observing these units, it was implicitly deduced that there already appeared some variability in the types of neuron APs recorded on day-20. One type of observable waveform profile (Fig. 1, example units 1, 6, and 9) was emphasized by an AP with a negatively-peaking trough. Another type of noticeable AP waveform shape (Fig. 1, example units 2, 4, 5, 7, and 8) was characterized by an initial positive peak, followed by a negatively-peaking trough of similar magnitude. The differences in amplitudes of these waveforms might be due to varying distances of recorded neurons respective to MEA electrodes, or due to an underlying difference in the type of neuron APs. Finally, the third type of discernible waveform shape (Fig. 1, example unit 3) was made prominent by a positive peak, and no clearly observable negative trough. Altogether, examining voltage traces of individual neuron APs revealed three discernible types of waveform profiles: negative, biphasic, and positive.

**Fig. 1.**
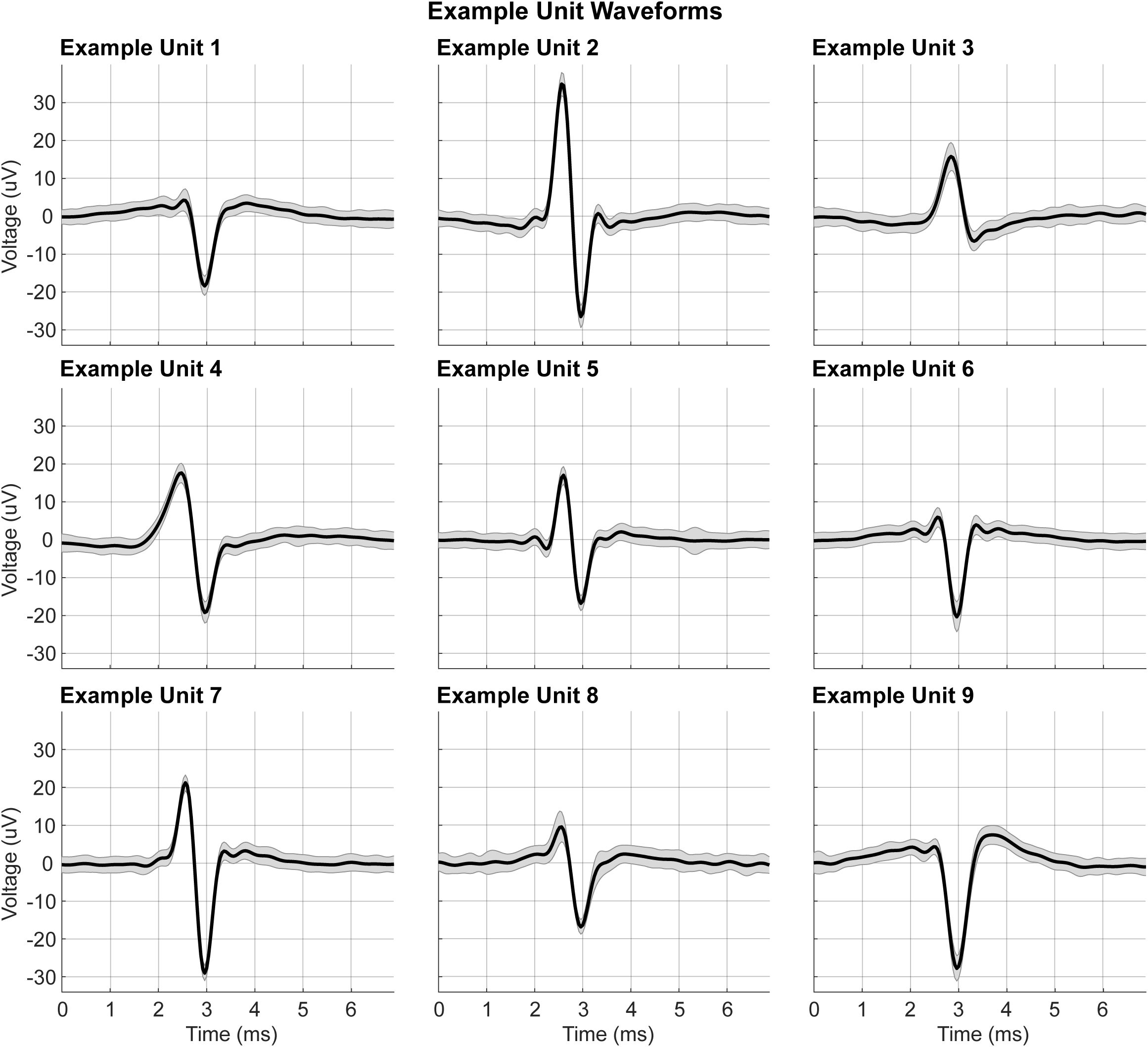
Example putative neuron (unit) AP waveforms. The plotted waveforms represent an average from all spikes detected per unit during the 10-minute MEA recordings. Shaded regions resemble one SD of the signal above/below the average voltage trace. The nine selected units were randomly picked from the GC MO model recording on day-20. The waveform time cut-outs were automatically computed by the spike-sorting algorithm. Each waveform cut-out represents 173 frames recorded at 25kHz sampling frequency, which translates to a time-window of 6.92ms.

### 3.2 Clustering Neuron AP Waveforms Uncovered Four Distinct Groups of Spike Profiles

After initially observing different spike-sorted waveform profiles, the subsequent step involved clustering units pooled across timepoints. Merging resulted in a dataset comprising of 282 putative neuron units from the GC MO model. Results of the threshold clustering established four clusters, each containing average AP waveforms that were grouped based on unit-waveform shape features. Fig. 2 depicts average waveforms from the four obtained clusters. The waveforms in Fig. 2 were normalized to voltage baselines, enabling a more comprehensible visualization of waveform shapes without the variance of amplitudes across units. The shaded regions surrounding the thick traces represent one SD margins around waveform averages of units in each respective cluster. Individual example waveforms of units were plotted in the top-right corners of the plots.

**Fig. 2.**
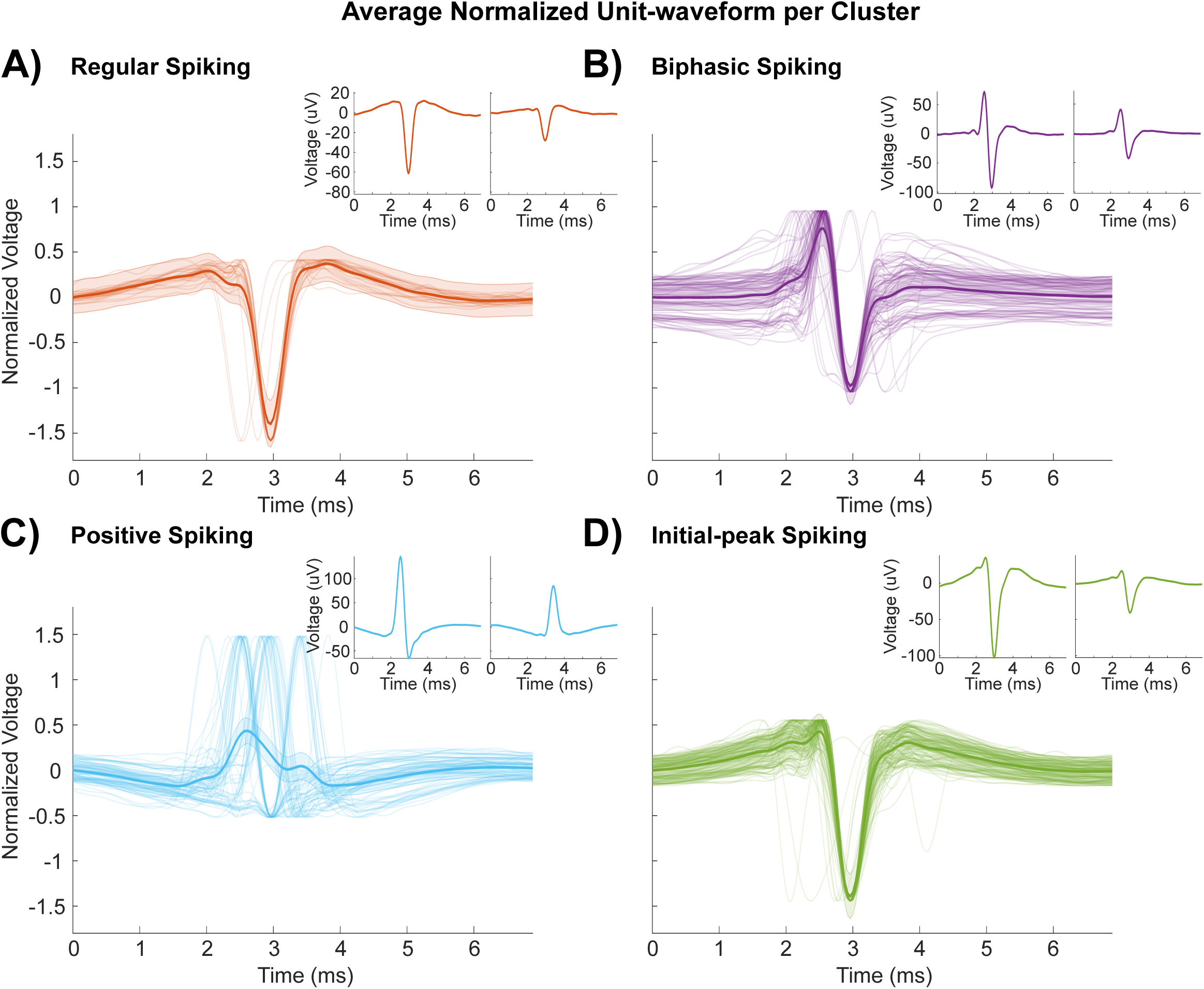
Four clusters of unit waveforms categorized by the clustering analysis. A-D) Each labelled cluster plot depicts normalized waveform cutouts of units belonging to the respective cluster plotted in the background, overlapped by an average normalized voltage trace. The shaded area around the thick average trace represents one SD of the mean signal. Following normalization in the range from -1 to 1, the offsets of voltage trace baselines were calculated to center the signals at 0 on the y-axis. Additionally, two cluster unit waveforms were randomly chosen and plotted in the top-right corners of the cluster figures. The two example waveforms were not normalized to highlight variability of spike amplitudes across clusters.

The first cluster (Fig. 2A) of unit waveforms was initially marked by a slow and minimal voltage increase followed by a negative trough, which lasted approximately 1ms from the depolarization to repolarization. The voltage then gradually recovered to baseline after ∼1.5ms. This cluster was labelled as containing regularly spiking units. The second cluster (Fig. 2B) of APs was distinguished by a sudden rapid voltage increase up to a peak accompanied by a negative trough of similar magnitude from the voltage baseline. Noticeably, the waveforms appeared to have different amplitudes prior to and after the AP, which resulted in a low interpretability of the recovery phase in this cluster. Nonetheless, given the similarly shaped positive and negative peaks, the waveforms of this cluster were further referred to as exhibiting biphasic spikes. The third group of waveforms (Figure-2C) was notable by a small voltage decrease succeeded by a positive peak and a gradual recovery to baseline. However, the individual positive peaks appeared to be asynchronously positioned along the x-axis, which obscured the average voltage trace of this cluster. Despite that, the waveforms in this cluster were termed as positively spiking. This label was attributed with reference to previous research by Sun et al.(Sun et al. 2021), where a cluster of positively spiking units was identified, despite having a similar degree of noise. The last cluster (Fig. 2D) was summarized as closely resembling unit APs in the first cluster (Fig. 2A), with the distinction of a small initial amplitude peak preceding the more discernible negative depolarization. Apart from this difference, the two clusters of unit waveforms appeared to be alike. Therefore, units in the last cluster were referred to as portraying ‘initial-peak’ spike waveforms.

An individual waveform cluster identity should not only be attributed by the clustering analysis, but also through its similarity with other waveforms in that respective cluster. This similarity, which also represents a measure of cluster quality, was not purely based on visual inspection. Another method of judging cluster quality was by inspecting cluster cross-correlograms. Fig. 3 illustrates cross-correlations across the waveform time-axis for all units in the four identified clusters. Evidently, most unit cross-correlations in the regular, biphasic, and initial-peak spiking clusters were ≥0.8, representing a high degree of similarity between the clustered waveforms, even though a few outliers persisted in each cluster. Conversely, the positive spiking cluster had roughly equal proportions of positively and negatively cross-correlated units in the range from -0.4 to 0.8. This noisy attribute of the positive spiking cluster was nonetheless accepted, given that all units displayed unique positive depolarization waveforms compared to the rest of the clusters.

**Fig. 3.**
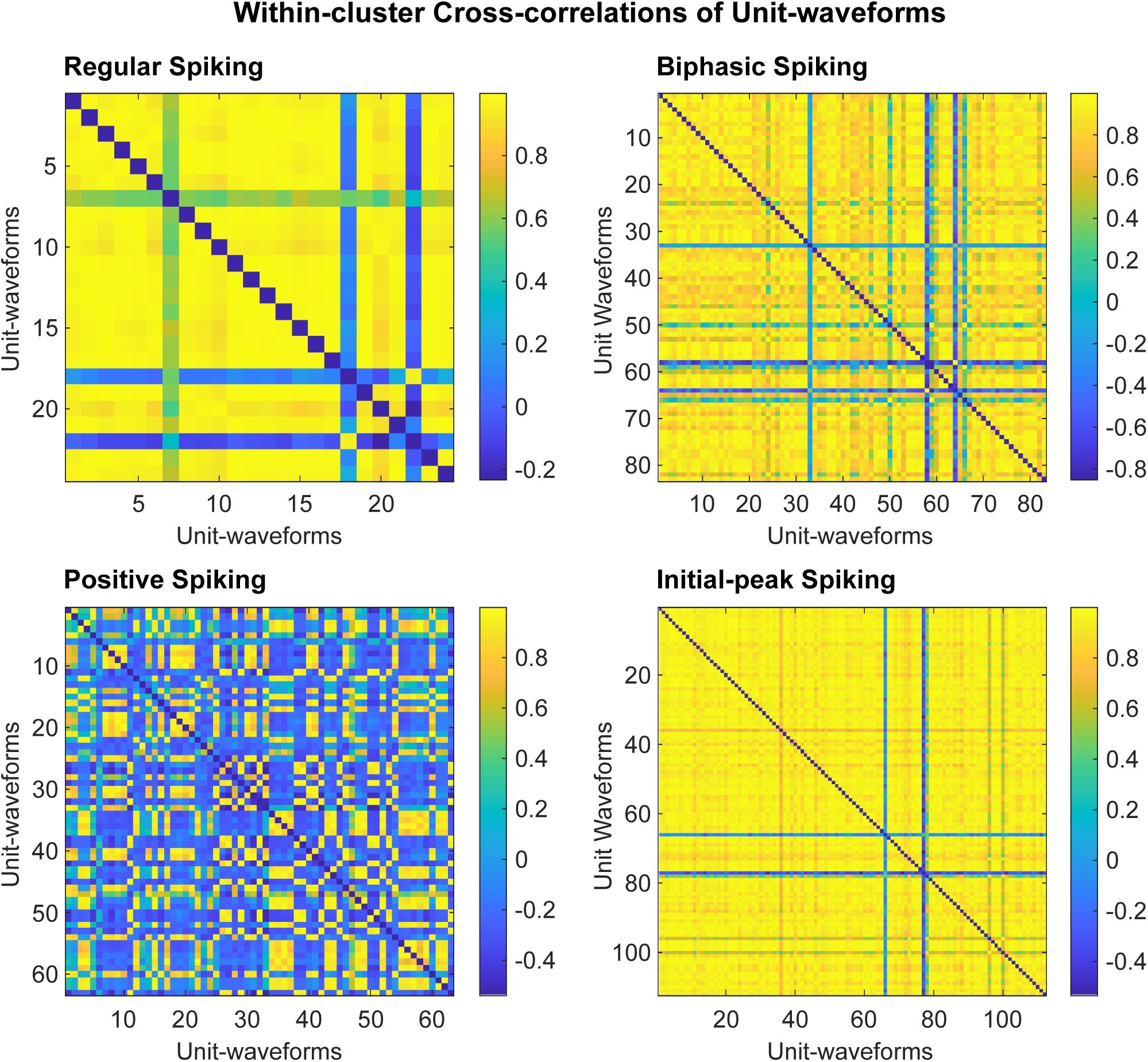
Cross-correlograms of unit waveform members per spike waveform cluster type. Each square in the plots represents a cross-correlation between two units across the waveform time-axis. The color, as indicated by colourmaps on the side of the plots, indicates the degree of positive/negative correlation among the respective units. The diagonal of the cross-correlograms was intentionally set to 0.

Judging the cluster quality was also approached from a different perspective, in terms of unit waveform positions in the t-SNE spatial 2D representation (Fig. 4C). In this latent space, each spike waveform was manifested as a vector position, according to its similarity and dissimilarity from other unit waveforms. In short, units proximal to each other possess a higher degree of waveform shape similarity compared to more distant units. Henceforth, the t-SNE representation was employed to highlight positions of clusters and spike waveforms in latent space. The unit waveforms were labelled with respect to cluster categories. The positive spiking cluster appeared to be a group of three more compact groups of units. This was ascribed to the noisy nature of waveforms in this cluster, not being aligned accurately on the time axis (fig. 2C). On the other hand, the t-SNE might have positioned the positive AP waveforms into three lower order groups as there might be further distinctions between types of positive spike waveforms. Focusing on the biphasic spiking units, the cluster center seemed defined, however, numerous outliers appeared present. The outliers were distinguishably located in proximity of the other three clusters, majority close to the positive spiking cluster. Lastly, the remaining two clusters appeared to have less of a well-defined cluster center and were surrounded by a few counts of biphasic spiking units. Yet, both the regular and initial-peak spiking clusters were still located at some distance from other clusters. In addition, an imbalance of unit waveforms favored the cluster identity of the initial-peak spiking units, with a lower number of regular spiking units located at the initial-peak spiking cluster center. The absence of a clear border between the two clusters signified their more pronounced similarity of waveform profiles compared to the rest of the clusters. When returning to Fig. 2, this close resemblance of average cluster waveforms becomes more apparent. Nonetheless, the two clusters were considered different, mainly leaning on the observation that regular spiking cluster waveforms have not portrayed the early peak before the negative depolarization that denoted the initial-peak spiking cluster.

**Fig. 4.**
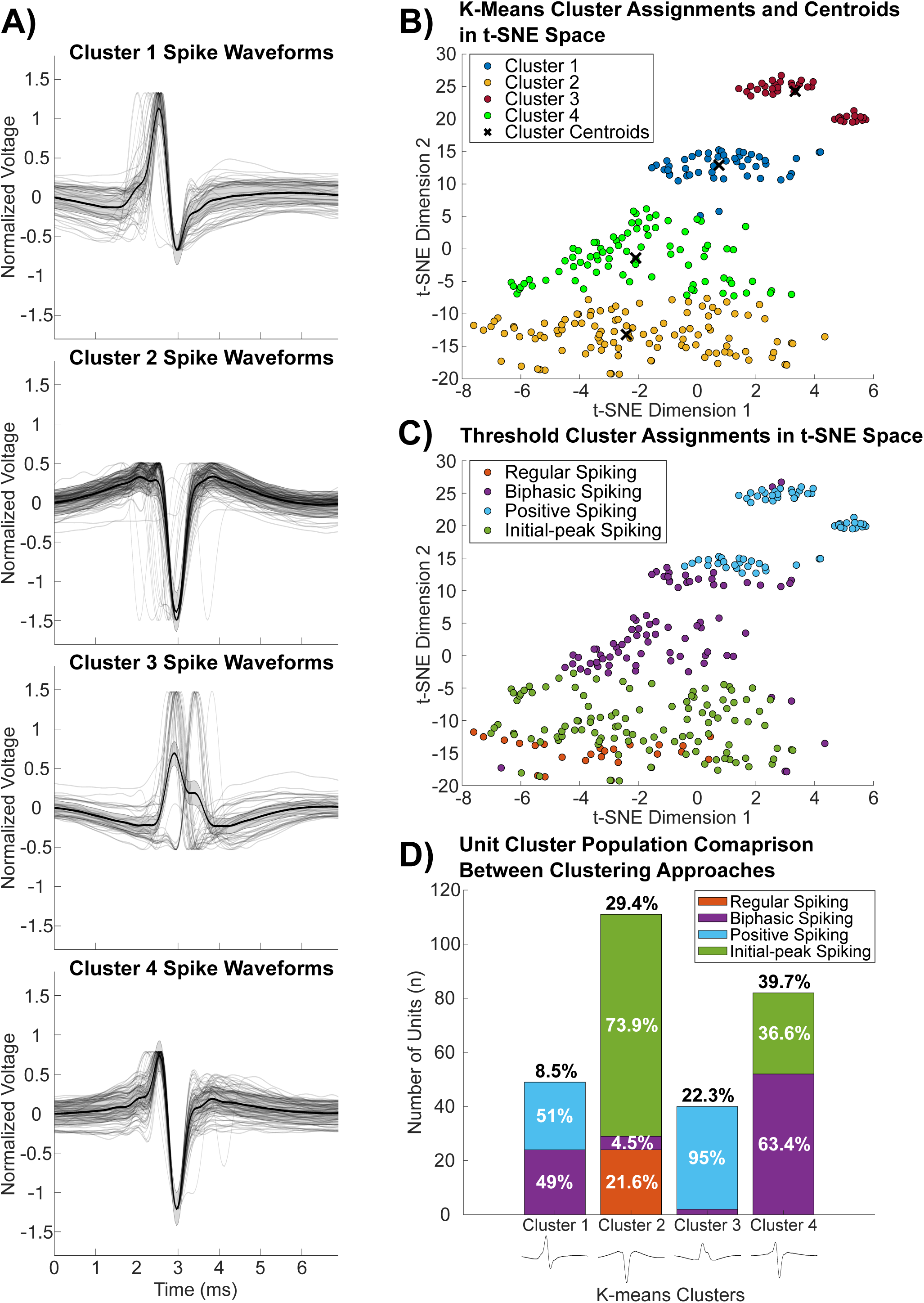
Overview of k-means clustering results in comparison to threshold clustering. A) Visualization of normalized average cluster AP waveforms obtained from the k-means clustering output. B-C) Unit waveforms represented as points of vectors in 2D t-SNE space. Waveforms were color-coded according to their cluster identity, as defined in the legends, resulting from k-means (B) and threshold (C) clustering analyses. D) Cluster population comparison between the automated and threshold approaches. The bars display counts of unit members per cluster identified by the k-means algorithm, percentage of cluster units from the total indicated above the bars in black. The bars were partitioned according to the threshold clustering waveform categorization as percentages in white, clusters labelled in the legend. Notably, the regular spiking unit identity was not recognized by the k-means algorithm, instead all regular spiking units were groups together with most of the initial-peak spiking units.

Overall, findings from threshold clustering highlighted the occurrence of four visually distinct clusters of putative neuron AP waveforms (Fig. 2). Yet, some clusters displayed a few potentially misclassified waveforms outliers (Fig. 3), and the t-SNE analysis surfaced some further speculation on whether the regular and initial-peak spiking waveforms were firmly categorized as different waveform profiles (Fig. 4C). Considering this, more evidence was necessary to discern whether these clusters reliably represented different types of putative neuron AP waveform profiles.

### 3.3 Comparing Spike Waveform Types Presented Differences Between Clusters and Clustering Approaches

Beyond threshold clustering of putative neuron AP waveforms into groups based on their profile resemblance, it was considered whether the waveform profiles varied when applying a more automated clustering approach. As threshold clustering obtained four waveform clusters, the preferred automated method was using k-means clustering, where the number of data partitions may be pre-determined. Results of the k-means clustering analysis showcased four clusters of spike waveforms depicted in Fig. 4A. The identified cluster waveform profiles elicited similarities but also some discrepancies with the threshold clustered waveforms. Notably, clusters one and four appeared to resemble the biphasic spike waveform shape (Fig. 2B), with cluster one demonstrating a slightly more pronounced positive peak before the negative trough. Additionally, cluster two waveforms were comparable to a combination of regular (Fig. 2A) and initial-peak (Fig. 2D) spike waveform profiles. Lastly, waveforms in cluster three presented a shape matching the positive spiking waveforms (Fig. 2C). Beyond reporting graphical features of cluster waveforms, the k-means output t-SNE representation was inspected to compare the categorized waveform positions in latent space (Fig. 4B) with the threshold clustering results (Fig. 4C).

Contrasting the clusters in t-SNE space highlighted that the regular and initial-peak spiking clusters formed a single cluster in the automated approach. On the other hand, the biphasic spiking cluster appeared divided into two clusters by the k-means algorithm, each partly including units from either the positive or initial-peak spiking clusters. To report interpretations about overlaps of clusters between the two clustering analyses, categorized units were counted and their cluster membership was compared across the two approaches. Fig. 4D showcases the unit populations of k-means clusters. Colors of the stacked bars indicated the portioned occurrence of units previously categorized by the threshold clustering analysis in percentages. This depiction more clearly highlighted differences between the two clustering methods. Cluster three was mostly representative of the positive spiking cluster, with the remaining positive spiking units being groups together with a proportion of biphasic spiking units in cluster one. Similarly, cluster four appeared primarily representative of the biphasic waveform cluster with some overlap of initial-peak spiking units. Interestingly, cluster two was largely dominated by initial-peak spiking units and the entire population of regular spiking waveforms. This observation hinted at the recurrent similarity between these two waveform profiles, given that the k-means algorithm partitioned them into a single cluster owning to their proximity in the t-SNE representation. Nonetheless, the clusters determined by the k-means algorithm lack the supervised nature of the threshold clustering approach, where units are grouped based on waveform profile features. In this sense, the automated k-means clustering was utilized to highlight potential differences in the obtained clusters of waveforms. Yet, the k-means algorithm grouped units in a latent t-SNE space, abstracted from the method of threshold clustering that is directly based on attributes of waveform shape. Henceforth, the threshold clustering results were the focus of interest for further investigation.

To assess potential differences in electrophysiological measures among the waveform types obtained from threshold clustering, comparisons of average cluster firing rates and spike amplitudes were conducted. Fig. 5A displays the distribution of binned unit firing rates. Notably, most neurons appeared to have a firing rate lower than 1Hz. It was further analyzed whether unit waveform clusters showed variability in average firing rates (Fig. 5B). The non-parametric test of ranks showed the average firing rate was significantly different between clusters, Х^2^(3,278)=10.40, *p*=.015. However, the following pairwise comparison tests revealed no specific average firing rate differences between the clusters.

**Fig. 5.**
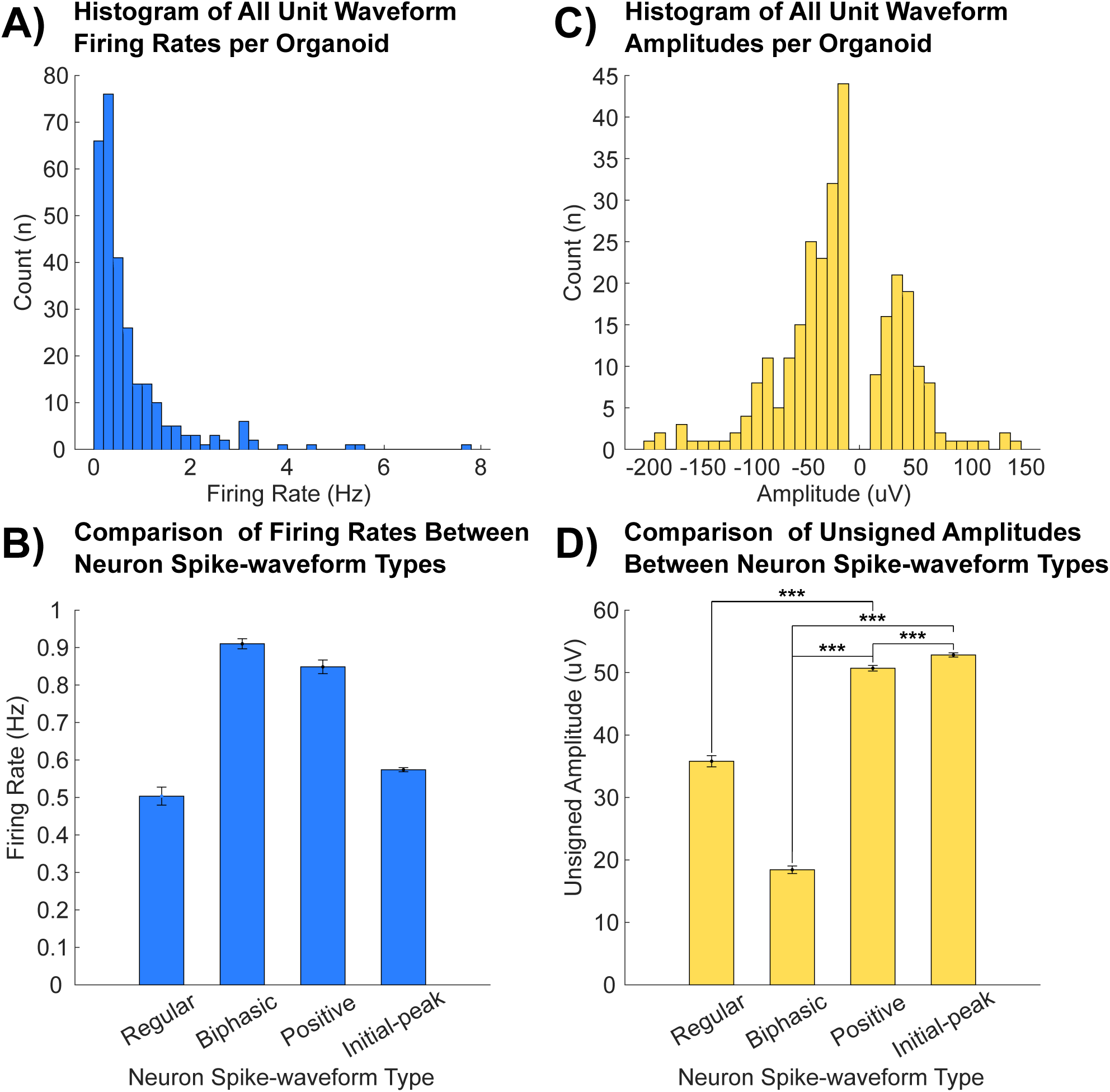
Graphical representation summarizing comparisons of average firing rate and spike amplitude between AP waveform types. A) Histogram depiction of binned unit firing rates. B) Bar chart demonstration of average cluster firing rates compared between clusters. There was no detectable difference of the firing rate among clusters. C) Histogram of binned unsigned amplitudes of all units. D) Bar chart illustration of cluster average spike amplitudes compared between clusters. The amplitudes were unsigned to facilitate comparisons of positive and negative amplitudes among waveform clusters. The mean amplitudes of regular and initial-peak spiking clusters were not significantly different. Additionally, the average amplitude between regular and biphasic spiking clusters was not significantly different. All additional pairwise comparisons of average cluster spike amplitude elicited significant differences among the groups. Note. * p<.050, ** p<.010, *** p<.001.

Shifting the focus to unit spike amplitudes, and considering that the amplitudes recorded via MEA devices can vary greatly based on distance of electrodes from the neurons, a histogram of amplitudes was produced for inspection (Fig. 5C). Here it was outlined, that even though bins on the distribution tails had considerably larger spike amplitudes, the bulk of units had lower amplitudes. Consequently, it was analyzed whether the average spike amplitude varied across waveform clusters (Fig. 5D). The Kruskal-Wallis test indicated a significant difference in average spike amplitude among clusters, Х^2^(3,278)=147.91, *p*<.000. Tests of six additional post-hoc pairwise comparisons revealed that the average amplitude was statistically different between four of the compared groups. The individual comparisons and associated p-values are presented in Table 2. Notably, the multiple comparison tests found no dissimilarity between the average amplitude of regular and initial-peak spiking clusters. In addition, there was no significant difference of amplitude among regular and biphasic spiking clusters. Altogether, observations suggested that clusters of waveform types demonstrated not only variations in waveform profile, but also in the average amplitude strength. This reinforced that the clusters resembled distinct types of neuron AP waveforms.

**Table 2.**
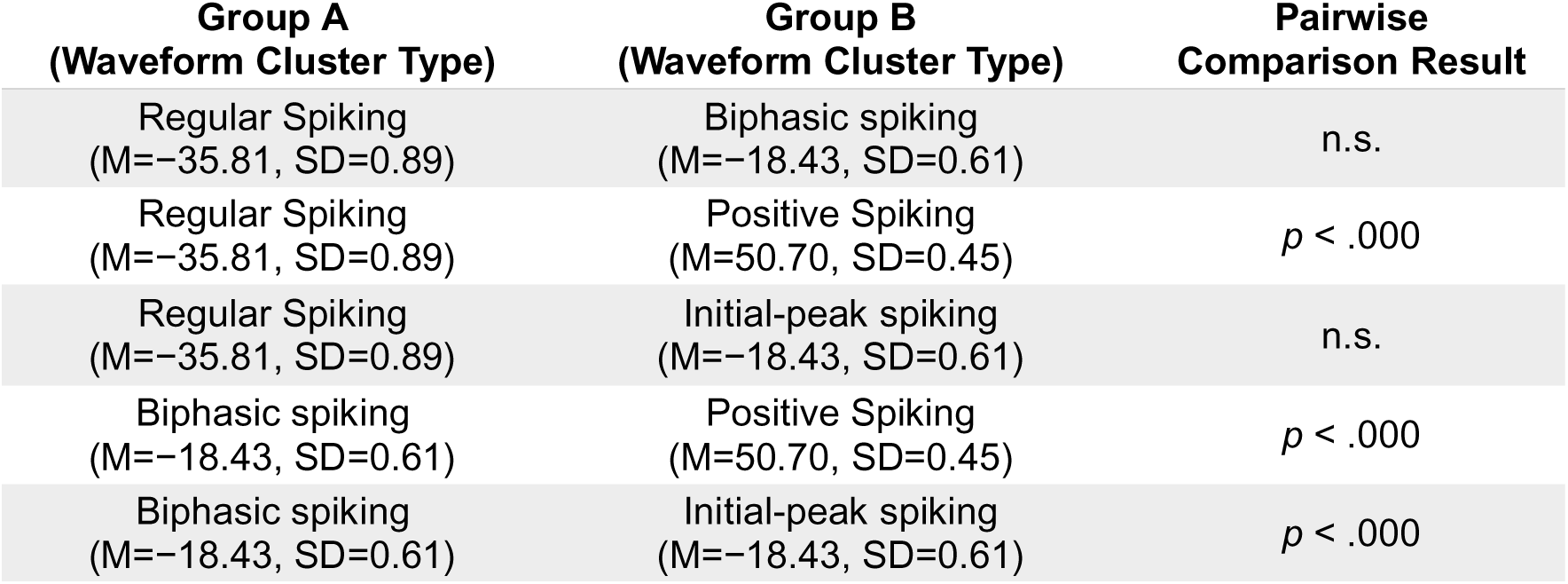
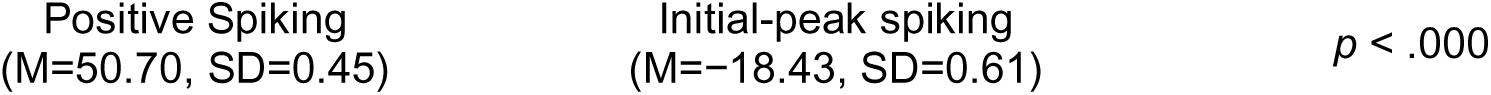
Post-hoc multiple comparisons between average cluster amplitudes. All six possible pair combinations of groups were comparatively tested. The table presents the resulting p-values from the pairwise comparisons of group A and group B cluster combinations. This analysis was conducted after observing a significant difference of the average amplitude between clusters from the Kruskal-Wallis test.

## 4 Discussion

In this study, the aim was to assess whether clustering of spontaneous electrophysiological activity could assist in identifying different types of AP waveforms exhibited by neurons within MO models. This was probed by constructing a computational pipeline to group neuronal spike waveforms extracted from MEA recordings of LFP activity into distinct categories. Establishing such an approach with electrophysiological data could be utilized to link waveform types to known neuronal populations, enabling separate analyses of electrophysiological activity of different neuronal cells. More specifically, the potential of isolating DNs for further analyses could benefit current research of neurodegenerative disorders like PD.

Findings from the spike-sorting analysis elicited that electrophysiological activity from MEA recordings appeared to be comprised of three different spike waveform shapes (Fig. 1). Threshold clustering results presented four groups of putative neuron spike waveform profiles (Fig. 4). These clusters were labelled as regular, biphasic, positive, and initial-peak spiking waveform types. The biphasic spiking cluster (Fig. 2B) was distinguished by waveforms with a positive peak volowed by a negative trough of similar magnitude before returning to baseline voltage. The positive spiking cluster waveforms (Fig. 2C) showcased positive depolarizations with a unique absence of a clear negative trough. Being mostly homogenous in shape, regular and initial-peak spiking cluster waveforms (Fig. 2A,D) were marked by a negative depolarization followed by a repolarization and gradual recovery phase. An observable distinction between these clusters was a slight initial positive peak of voltage preceding the negative trough in the Fig. 2D waveform. Hence, it was termed the initial-peak spiking cluster. Through cross-correlograms it was established that the regular, biphasic, and initial-peak spiking cluster units had satisfactory degrees of within-cluster correlations (Fig. 3). The positive spiking cluster displayed positive and negative unit cross-correlations that were considerably lower than in other clusters. It was reasoned to be caused by timepoint disparities of positive peaks in this cluster (Fig. 2C), resulting in some units being correlated positively and negatively to the rest. Nonetheless, all the units exhibited unique positive APs, so the misalignments across time and lower cross-correlations were acceptable. The waveform shape clusters were also examined in a t-SNE representation (Fig. 4C). Here, the waveforms were vectorized onto a 2D plane based of their similarity with neighboring waveforms. The greater the distance between waveform positions, the more dissimilar their shape. The positive spinking cluster was observed as a collection of three more compart visually discernible groups of waveforms, which might be attributed to the timewise misalignments of waveforms from this cluster. Furthermore, the biphasic cluster contained outliers that were proximal to other clusters of waveforms. This could be because the biphasic AP waveform profile was marked by both a positive and negative peak, resulting in its resemblance to other cluster waveforms. Hence, regardless of outliers, the cluster category of biphasic spike waveforms was accepted as reasonably substantiated. The regular and initial-peak spiking clusters were closely positioned, the former enveloping the latter in the t-SNE space. Arguably, this was because waveforms from these clusters had similar profiles, apart from the initial peak before the depolarization which was less pronounced or even absent in regular spiking waveforms (Fig. 2A).

Further, the threshold clustering results were probed in comparison to an automated k-means clustering approach. Three of the four waveform clusters displayed in Fig. 4A highlighted the retrievability of positive and biphasic waveform shapes by the k-means algorithm. However, regular and initial-peak spike waveform clusters appeared partitioned into a single cluster. This was more evident from the unit populations of the k-means clusters overlapped by the categorization of threshold clustering results in Fig. 4D. Here it was outlined that k-means clusters three and four were majorly represented by either positive or biphasic spiking unit waveforms. Cluster one was almost equally populated by positive and biphasic units which may be due to their proximity in the t-SNE space. Most interestingly, the k-means cluster two largely consisted of initial-peak spiking units and the entire population of regular spiking units. This further supported overcategorization of the negatively peaking waveform profiles into two distinct clusters, conveying the idea that regular and initial-peak AP waveforms may be generalized into a single cluster given their predominant resemblance.

These observations prompted comparisons of other electrophysiological metrics between the threshold clustering waveform types, namely average cluster firing rates and spike amplitudes. Although the average firing rate was different across the four clusters, post-hoc multiple comparison tests revealed no specific variations between individual cluster pairs (Fig. 5D). Conversely, average spike amplitude was more heterogeneous across the waveform clusters (Fig. 5D), further establishing categorical uniqueness of the waveform profiles. This was with exception to the average regular spike amplitude, which was not different compared to initial-peak and biphasic cluster amplitudes. These findings, in addition to the previously made remarks, pointed to the conclusion that regular and initial-peak spike waveforms have been over-clustered and should arguably resemble a single type of waveform profile. Another line of reasoning was that there is a lack of data, as the dataset only contained 282 units from GC MO recordings. Hence, clusters had a low sample size of waveforms to firmly establish their varying identities, as the utilized non-parametric comparison methods were less sensitive to reveal statistical differences between groups. Future data collections to repeat this methodological approach may highlight more evidence on this subject. Altogether, it was concluded that the threshold clustering analysis reliably identified three types of unit waveforms, namely biphasic, positive, and regular (merged with initial-peak) AP waveform profiles. Thus, consistent with the primary hypothesis, the clustering approach was able to classify putative neuron spike waveforms from recorded LFP activity into three distinct neuronal waveform types. Interestingly, these were also the three waveform types first spotted in Fig. 1 of example units from a single MEA recording.

This was in line with previous studies, where the application of a similar clustering method yielded comparable results. Sun et al.(Sun et al. 2021) found five different waveform types from extracellular recordings in the cat primary visual cortex, validating this approach. Jung et al.(Jung et al. 2023) replicated these clustering results with data recorded from the wallaby primary visual cortex, also observing similar categories of waveforms, but distributed in different proportions. More importantly, two of the waveforms identified in this prior research appeared visually alike to the presently obtained positive and regular spiking clusters. However, the fast-spiking, triphasic-spiking, and compound-spiking neuron waveforms were not reflected in current findings. This could be due to the difference in recorded samples. As mentioned earlier, neuronal cells have a varying development, proliferation and structural organization across species(La Manno et al. 2016). Furthermore, neurons in human MOs(Zagare et al. 2022) differ from those in the mouse V1(Gouwens et al. 2019), notably by the absence of DNs in the latter. Hence, the absence of waveform clusters in contrast to previous findings was attributed to the type of cell community in human MOs. Furthermore, the presently employed clustering analysis was adapted to fit observations of spike-sorted unit waveforms, creating further disparity from the previously applied method. Nonetheless, the current approach appeared to hold the premise of clustering neuron APs into defined categories of elicited waveform profiles.

Regarding the secondary hypothesis of this project, we subsequently considered whether it was possible to link observed waveform types to known neuronal populations within the recoded MO model. It was intuited that it may be possible to distinguish neuronal identity from observed clusters of waveform types by comparing the waveforms to validated templates of specific neuron AP profiles. For instance, focusing solely on DNs, Grace and Bunney(Grace and Bunney 1983b, 1983a) established fundamental extracellular/intracellular mechanisms and morphological knowledge on APs of midbrain DNs in rats. The APs comprised of two different spike segments. The first, termed initial-segment spike, was generated at more distal dendritic sites from the DN soma with a low depolarization threshold. The second type, named somatodendritic spike, was produced with considerably higher amplitudes compared to the initial-segment spikes. The somatodendritic spikes also had a higher depolarization threshold and were primarily depolarized from strong inputs of pacemaker potentials. It was also reported that a full-scale depolarization at the axon hillock mainly required the somatodendritic spike, not only initial-segment spikes. Surprisingly, the waveform shape of initial-segment spikes was marked by a positive peak and no negative trough, appearing resemblant of the positive spiking waveform cluster observed from the current data (Fig. 2C). Additionally, the somatodendritic spike waveform shape aligned closely to the biphasic AP waveform cluster shape (Fig. 2B). Hence, it was reasoned that positive and biphasic spike waveforms may resemble initial-segment and somatodendritic DN spikes, being produced by the same type of cell but at different locations with respect to the soma. What obscured this postulation was that positive spiking waveforms had a higher average amplitude compared to the biphasic spiking cluster (Fig. 5D). While initial-segment spikes should have a lower amplitude compared to somatodendritic spikes(Grace and Bunney 1983b). Instead, this observation might be simply explained by the intuitive idea that DN somata were located further from MEA electrodes above the coating layers. Dendrites may have grown through the layers towards the electrode surface, making their amplitude appear larger amid recordings. However, there was no evidence for such deductions in this experimental design.

Taking an alternate perspective, the currently observed percentage populations of neuronal waveform types might suggest a consensus regarding the attribution of positive and biphasic spiking waveform types to DNs. Previous research examined MOs using flow-cytometry(Monzel et al. 2017). Researchers revealed that roughly 66.6% of MO neurons expressed the tyrosine hydroxylase (TH) protein, which predominantly resides in DNs. Assuming biphasic and positive waveform clusters originated from DNs, their combined incidence would have been 51.8% in the MO. Nonetheless, there was no statistical support for these compositional comparisons as only one organoid model was analyzed, without any additional sample replicates. Future research could record from multiple replicates of an organoid model and cell line. This would enable statistical comparisons between compositions of neuron waveform types present within the recorded organoid models.

Nevertheless, more evidence was required for attribution of positive and biphasic waveform clusters to DNs. Researchers have previously conducted patch-clamp recordings of DNs in rats, deriving shapes of extracellular waveforms by the first differential of the intracellularly recorded waveforms(Ungless and Grace 2012). This was to confirm that waveforms obtained from actual in-vivo extracellular recordings originated from DNs. Results revealed that extracellularly recorded rat DNs exhibited APs labelled as biphasic waveforms. Even though the waveforms were resemblant of the biphasic spiking waveform cluster (Fig. 2B), the positive AP waveform profile (Fig. 2C) was not observed. More recent research showcased some further findings on this matter, employing computational modelling to simulate a 3D model of a DN(López-Jury et al. 2018). The extracellular data used to construct this model were recorded from rat DNs. Researchers utilized the model to simulate extracellular recordings from 42 locations around the DN soma, dendrites, and axon across different conditions. Findings revealed that the electrode location influenced the recorded spike waveform shape. Additionally, the researchers depicted waveform amplitudes on a logarithmic scale, resulting in some waveforms having considerably small amplitudes. Drawing on the results of this computational simulation study, and taking into account waveform amplitudes that would theoretically be detectable by the MEA, DN identity could once again be attributed to the biphasic spiking waveform profile (Fig. 2B). It was thence concluded that the biphasic cluster AP waveforms originated from DNs. This was not argued for positive spiking waveforms (Fig. 2C), as evidence was less converging in support of their resemblance to DN spike waveforms. Altogether, this aligned with the secondary hypothesis that spike waveform profiles could be linked to known neuronal populations, such as the DN.

However, some limitations were outlined. The present conclusion was primarily based on visual comparisons of waveform profiles to DN spikes depicted by previous research(Grace and Bunney 1983b, 1983a; Ungless and Grace 2012; López-Jury et al. 2018). The present experimental design did not include a method of reliably checking whether a biphasic spiking unit recorded by the MEA corresponded to an actual DN in the MO. Even though previous studies supported the presence(Zagare et al. 2022), estimated proportion(Monzel et al. 2017) and waveform resemblance of DNs in organoid models, these observations require an additional cellular identity verification of the neurons that exhibited the APs. One possible way of attributing clustered waveform profiles from electrophysiological activity in 3D organoids to actual neurons might be using a cell-type-specific genetic silencing experimental design. Future research could record electrophysiological activity before and after selective inhibition of neuronal populations. In one condition, DN activity could be suppressed through expression of an inhibitory chemogenetic or optogenetic construct under a TH promoter, allowing selective silencing of DNs while preserving the activity of other neuronal populations(Armbruster et al. 2007; Gradinaru et al. 2008; Roth 2016). A second condition could involve selective silencing of excitatory or inhibitory neurons using equivalent cell-type-specific genetic constructs. Electrophysiological activity and spike waveforms associated with each neuronal population could then be isolated by comparing the clustering analysis results of a control group to the silencing conditions described above. Observing whether a type of waveform cluster is abolished or retained following silencing of a specific neuronal population compared to the control waveform clusters would serve as a verification that an AP waveform profile originated from that neuronal population.

A second methodological design for improving the reliability of assigning clustered AP waveform profiles to specific neuron types could employ fluorescent microscopy combined with cell-specific staining. In this approach, DNs could be stained with a fluorescent marker(Blokhin et al. 2022; Lavrova et al. 2019). MEA recordings would then be combined with fluorescent microscopy of DNs. After spike-sorting the extracellular data, approximate locations of the extracted putative neurons could be correlated to locations of stained DNs with respect to MEA electrodes. This method could serve as an additional confirmation that a unit waveform and its associated waveform cluster corresponds to a localized DN. Finally, another way of distinguishing neuron APs from 3D organoid models could be by recording electrophysiological activity via two-photon calcium imaging. In this experimental design, a genetically encoded calcium indicator (GECI) would be expressed in a specific neuronal population such as DNs(Akerboom et al. 2013; Dana et al. 2019). Changes in calcium flux would only be detected in the population of neurons expressing the GECI. Taking this a step further, using multicolor GECIs would allow calcium imaging of multiple cell-type-specific neural populations collectively(Inoue et al. 2019). This might enable future research to conduct more advanced electrophysiological activity comparisons of specific neuronal cell types between diseased and healthy models. This complex methodological design would be challenging, yet it could serve as another verification to support attribution of AP waveform clusters to specific neurons that are electrophysiologically active in 3D organoid models.

Overall, we concluded that clustering of electrophysiological activity in an organoid model does aid in identifying compositions of AP waveform types present within the organoid. We further concluded that spike waveform clusters may be linked to actual neural identities, yet additional evidence from cell-identity verifications would strengthen this conclusion in future improvements of the experimental design. Regarding the scientific relevance of current findings, the employed clustering approach yielded the identification of three diverse AP waveform profiles in the MO. One of the spike waveform types was arguably originating from DNs. This is scientifically relevant as a novel analysis approach of organoid functionality, as it allows potential isolation of electrophysiological activity into groups resembling specific neuronal populations. This experiment has also provided a methodological insight into the advantages and drawbacks of clustering spike-sorted datasets of putative neurons to identify different types of spike waveforms. In the realm of pre-clinical research, this method could offer a fresh perspective on the possible analyses of electrophysiological data obtained from commonly employed MEA devices for recording organoid models. Future outlooks should include refinements of the experimental design using cell-type-specific genetic silencing, or fluorescent microscopy together with the proposed clustering approach. Altogether, this computational analysis indicated some potential benefits in the methodological toolbox for ongoing research into organoid model development and the investigation of neurodegenerative diseases such as PD.

## Author contributions

Concept (JO), Design (JO, DF), Definition of intellectual content (JCS), Literature search (JO, ASZ), Experimental studies (JO, DF, ASZ), Data acquisition (JO), Data analysis (JO), Statistical analysis (JO), Manuscript preparation (JO), Manuscript editing (JO, ASZ, DF, JCS), and Manuscript review (JO, JCS).

## 5. Acknowledgements

The authors thank Dr. Jared Sterneckert for providing iPSC lines. The JCS lab is supported by the Fonds National de la Recherche (FNR) Luxembourg (PRIDE21/16749720/NextImmune2). We also thank the private donors who support our work at the Luxembourg Centre for Systems Biomedicine. The authors acknowledge support from the European Union’s Horizon 2020 1059 research and innovation programme H2020-FETPROACT-2018-01 under grant 1060 agreement 824070 (CONNECT).

This research was funded in whole by FNR Luxembourg. For the purpose of Open Access, the author has applied a CC BY public copyright license to any Author Accepted Manuscript (AAM) version arising from this submission.

